# Role of Sugar Stereochemistry on Structural and Free Energy Landscape of Double-Stranded Nucleic Acids

**DOI:** 10.1101/2020.08.15.252643

**Authors:** Anuj Kumar, Reman Kumar Singh, Amol Tagad, G. Naresh Patwari

## Abstract

The conformational landscape of 29-mer long four stereo variants of furanosal nucleic acids and their C2’ deoxy counterparts were explored using molecular dynamics and well-tempered metadynamics simulations. The ribose containing double-stranded nucleic acids exhibit helical structure, however inversion of C3’ and/or C2’ stereocentre of ribose results in structural modification. The free energy surfaces relative to the average twist angle and end-to-end distances suggests that the configuration at the C3’ position plays a pivotal role in determining the helicity. The C3’ stereocentre acts as toggle-switch for helix to non-helical structures in double-stranded nucleic acids. Thus, the ribose containing double-stranded nucleic acids result in well-organized helical structures, while those containing xylose and lyxose show a variety of structures, which include (circular) ladder configurations. Based on the present set of results, it can be inferred that ribose containing double-stranded nucleic acids form well-defined helical structures in contrast to their C2’ and C3’ epimers.

## INTRODUCTION

The chemical etiology of nucleic acids has been subject of great curiosity,^1^ and several synthetic approaches have been employed to address this issue by varying one or more ‘structural elements’ of the canonical nucleic acids viz., the nucleobases, the sugar and the phosphate linkage.^2,3^ Xeno nucleic acids (XNA) are sugar modified nucleic acids,^4^ which include chemical modification of the ribose sugar with several variants that may or may not include furanose sugar. An interesting question raised by Eschenmoser in this context is,^5^ “why did nature choose furanosyl-RNA and not pyranosyl-RNA?” and concluded based on the experimental evidence that pyranosyl-RNA binds more strongly and selectively relative to its natural counterpart the furanosyl-RNA.^5^ However, the primary question remains unanswered. Subsequently, many variants of XNAs with non-furanosyl moieties such as cyclohexene,^6^ glycol,^7^ hexitol,^8^ peptides,^9^ threose,^10^ and others have been reported, which demonstrate their ability to pair can be modulated by appropriately modifying the ‘sugar’ moiety even while the nucleobases are conserved to form Watson-Crick pairs. A large number of XNAs have been explored in various biological technologies such as antisense oligonucleotides, siRNAs, and aptamers,^11,12^ prominent among them being C2’ modified analogues. On the other hand, arabino nucleic acids, which have inverted C2’ stereocentre are known to equilibrate between single-stranded hairpin structure and canonical B-form.^13^

The ability of canonical nucleic acids to elongate and structurally transform, albeit transitorily, have been captured during the transcription process.^14^ However, a standalone ladder-like structure of canonical nucleic acids has not been reported, whilst structural transformation of helical B-DNA into ladder-like structure can be explored by quenching of dispersion energy component of the potential function^15^ or by force stretching.^16^ Interestingly, substitution of ribose with xylose results in a ladder-like structure for the double-stranded furanosyl-xylo nucleic acid (xyloNA),^17^ and its deoxy analog (DxyloNA).^18^ The structural landscape of xyloNA and DxyloNA, explored with the aid of molecular dynamics simulations, reveals their propensity to form ladder-like structures along with left-handed helix.^19–22^ Interestingly, the thermodynamic stability of DxyloNA was found to be higher than DNA,^23^ which is in contrast to the facile unzipping of xyloNA relative to RNA.^24^ While there has been a flurry of activity to explore the structure and dynamics of both native and deoxy versions of xylo nucleic acids, using both experimental and computational tools, investigations on nucleic acids constructs with the C2’ epimer of xylose, the lyxose, remain largely unexplored.^25^ An interesting question that arises at this stage is, “how does the configuration at the C2’ and C3’ stereocentres influence the structural landscape of double-stranded nucleic acids?” More specifically, “is the ladder-like structure of double-stranded xyloNA (and DxyloNA) is associated with the inversion of C3’ stereocentre, relative to RNA (and DNA)?” To this end, molecular dynamics simulations on double-stranded nucleic acids consisting of four variants of furanose and two of C2’ deoxy counterparts, with the sequence 5’-[GCAU(T)]γG-3’, structures of which are shown in Figure 1, were carried out to understand the role of sugar stereochemistry, and the results are discussed herein.

**Figure 1.**
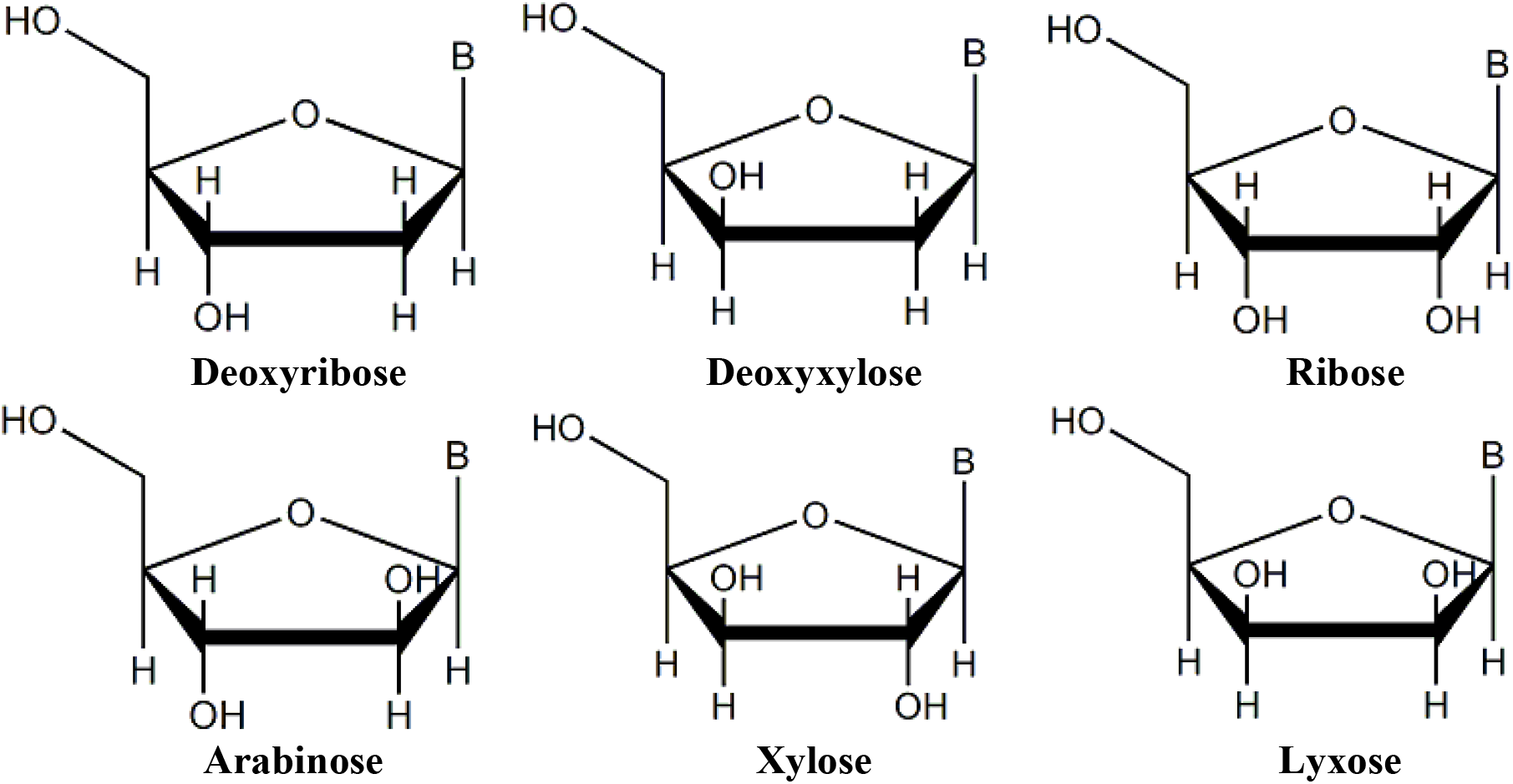
Structures depicting stereochemistry of various furanose sugars used in the present work.

## METHODOLOGY

The structures of the double-stranded nucleic acids of RNA, arabinoNA, xyloNA, and lyxoNA and their corresponding deoxy counterparts DNA and DxyloNA were explored using MD simulations. The initial structures of double-stranded B-RNA and B-DNA with sequence 5’-(GCAU)_7_G-3’ along with its complementary strand was generated using the make-NA server.^26^ The initial structure of the double-stranded xyloNA structure was obtained from Ramaswamy *et al.^20^* The structures of arabino- and lyxo-nucleic acids were generated by inverting the stereochemistry at the C2’ position of furanose sugar in the ribo- and xylo-nucleic acids, respectively. Further, the initial structure of deoxy-xylose form was generated by replacing the OH group with H in the xyloNA. All-atom molecular dynamics simulations were carried out for 29-mer double-stranded nucleic acids with Amber-14SB forcefield^27^ using double precision GROMACS 2020.^28,29^ The double-stranded nucleic acid was oriented along the principal axis of a rectangular box. The dimension of box was defined such that the solute should not interact with its periodic image. The simulation box was solvated with TIP3P^30^ water molecules along with 56 sodium cations in each case to achieve the electro-neutrality of the system. The potential energy of the system was minimized using the steepest descent algorithm^31^ with a tolerance force of 10 kJ mol^-1^ to remove any bad contacts and steric clashes. After energy minimization, the system was equilibrated in the NVT ensemble by employing a velocity-rescale thermostat ^32^ with a coupling time of 0.4 ps and a reference temperature of 300 K for 2 ns. The heavy atom of nucleic acid was restrained using harmonic forces with force constant 1000 kcal Å^-2^. This was followed by NPT equilibration for 5 ns using Berendsen barostat^33^ with coupling time of 2 ps and reference pressure of 1 bar. For the production runs, the temperature of the system was maintained at 300 K using a Noose-Hover thermostat with coupling constant of 0.4 ps and the pressure of the system was maintained at 1 bar using the Parrinello-Rahman barostat ^34^ with a relaxation time of 0.6 ps. The equations of motion during the simulations were integrated using the leapfrog algorithm ^35^ with a time step of 2 fs. For each of the six double-stranded nucleic acids, five independent trajectories of 100 ns were performed with different initial velocity and by keeping other protocol as mentioned above. Further, the conformational space of all the double-stranded nucleic acids considered in the present work, was sampled using well-tempered metadynamic simulations^36^ by biasing with a Gaussian potential of 1 kJ mol^-1^ height with sigma of 0.06 along two reaction coordinates, the average twist angle (χ), and the end-to-end distance (*l*). The twist angle measures the helicity of double-stranded nucleic acids while the end-to-end distance measures the contour length of nucleic acid as schematically depicted in Figure 2. In this analysis the base pairs at two ends were neglected due to larger amount of fluctuations.

**Figure 2.**
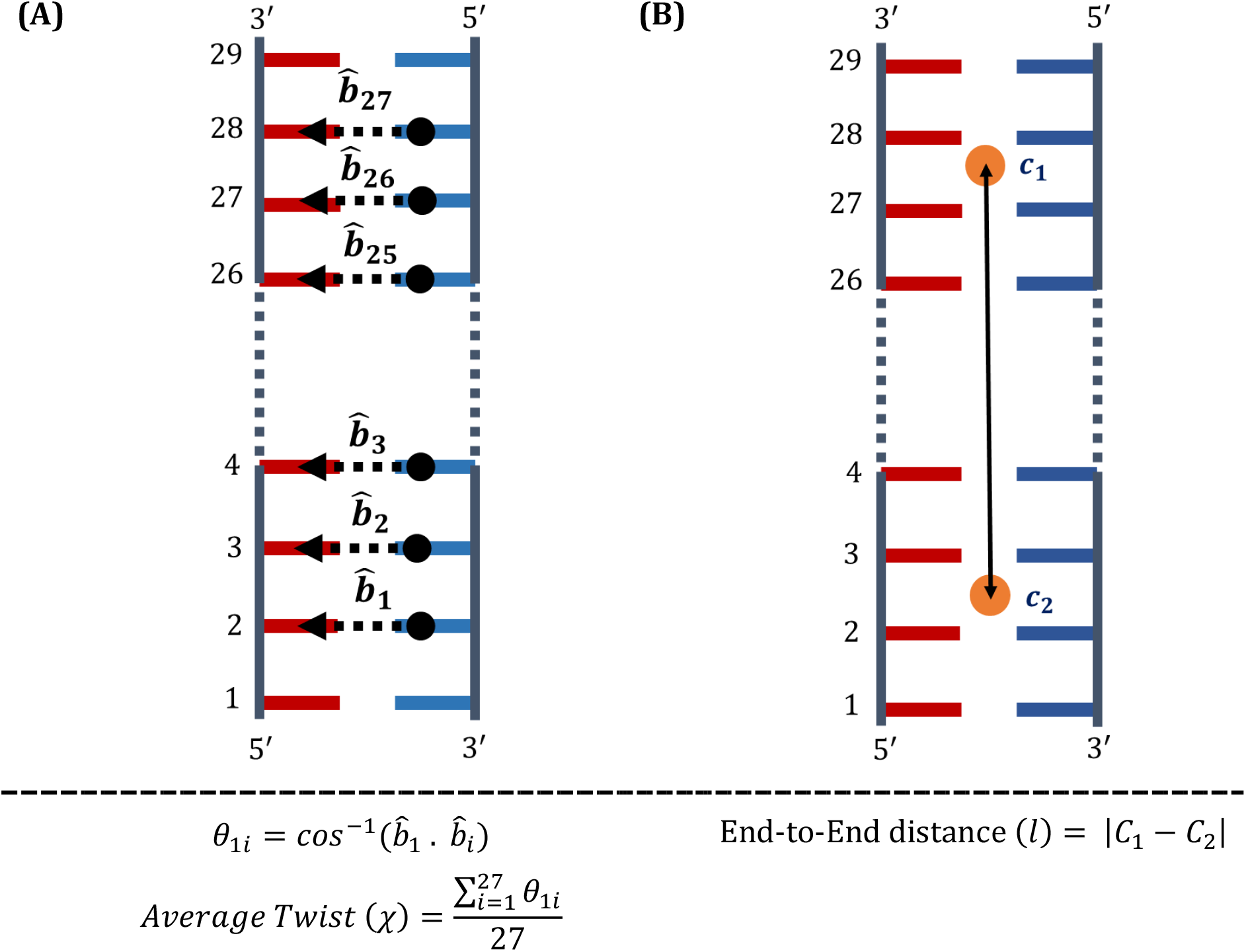
Description of the reaction coordinates (A) average twist angle (χ) and (B) end-to-end distance (*l*) used to calculate the structural and free energy landscape. In (A) 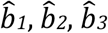, and others are unit vectors joining the centre-of-masses of inter-strand base pairs and *ci* and *c2* are centre-of-masses of the two adjacent Watson-Crick base pairs. In this analysis the base pairs at two ends were neglected.

## RESULTS & DISCUSSION

The structural evolution of all the six sets of double-stranded nucleic acids was followed through the course of the simulations and the converged structures at the end of 100 ns simulation are depicted in Figure 3. Figure 3 illustrates that for RNA, arabinoNA, and DNA retain right-handed helical structures throughout the simulation. On the other hand, for the xyloNA, lyxoNA, and DxyloNA an apparent loss of helicity is be observed and these structures cannot be classified as right-handed helices. *Prima-facie,* it appears that the inversion of configuration of the furanose ring at C3’ position in the xyloNA, lyxoNA, and DxyloNA vis-à-vis RNA, arabinoNA, and DNA respectively, results in loss of helicity. In an effort to statistical validity these results a total of five independent trajectories of 100 ns for each of the six variants of double-stranded nucleic acids were carried out. Figure 3 also depicts a plot of end-to-end distance (averaged over all the five trajectories) over the period of simulation. The end-to-end distance is conserved (around 7.8 nm) for the RNA, arabinoNA and DNA, while the corresponding plots for the xyloNA, lyxoNA, and DxyloNA show marked variation over the period of simulation. Another important structural feature is the distribution of twist angle (averaged over all the five 100 ns trajectories), plot of which also is shown in Figure 3. The average twist angle shows the narrowest distribution for arabinoNA centred around 100°, whereas DNA in comparison shown marginally broader distribution, once again centred around 100°. Interestingly, RNA shows a bimodal distribution one centred around 100° and other centred around 110°. The twist angle distribution for lyxoNA is broader with centre around 85°, while those for xyloNA and DxyloNA are much broader distributions with much lower twist angle. The lower values of average twist angles can be attributed to formation of non-helical structures.

**Figure 3.**
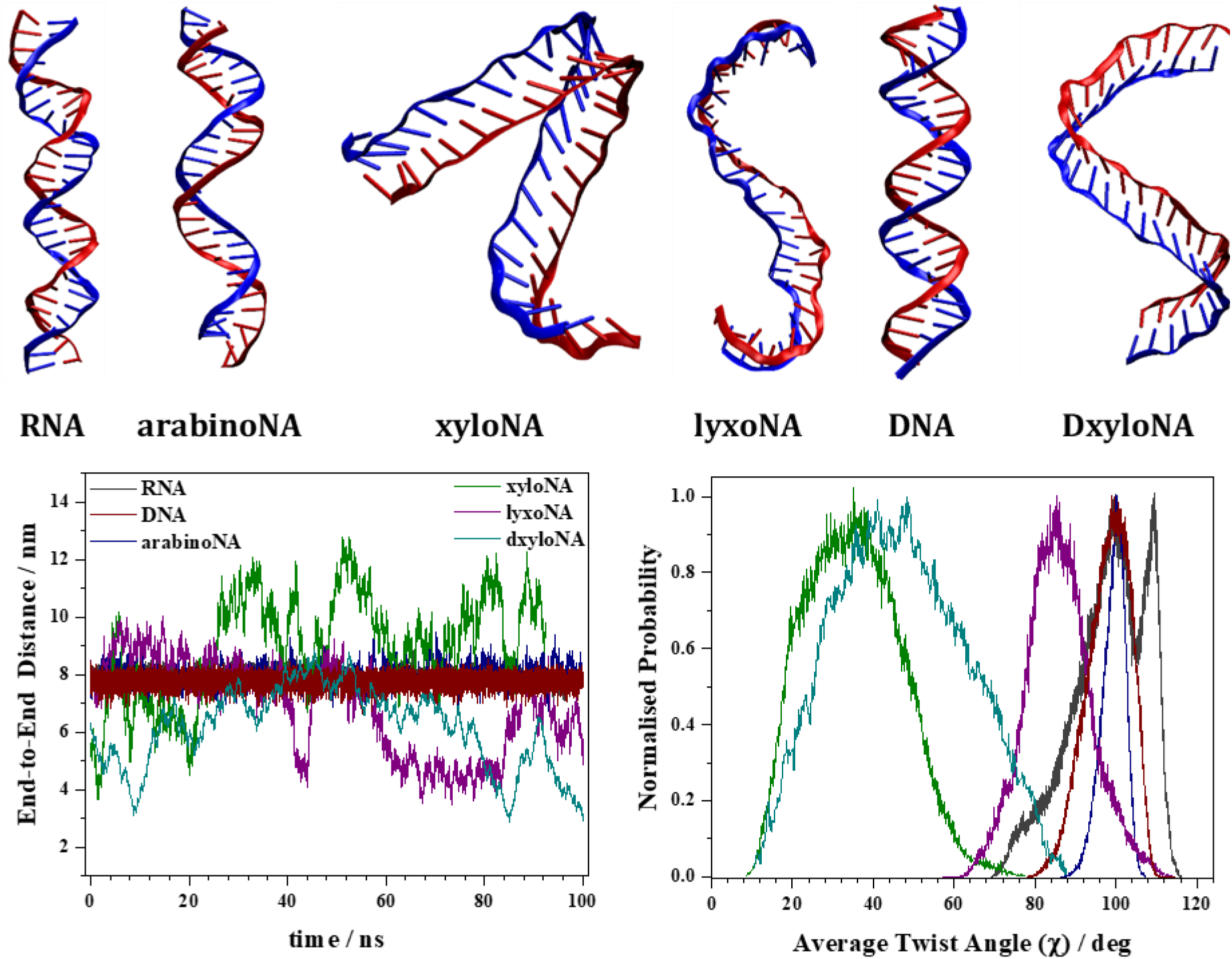
Snapshots of structures of double-stranded nucleic acids obtained after 100 ns of simulation. The structures of RNA, arabinoNA, and DNA appear as right-handed helices while the structures of xyloNA, lyxoNA and DxyloNA are non-helical. The bottom panel shows plot of end-to-end distance over the simulation time and twist angle distribution, both averaged over five 100 ns trajectories. The depicted structures and the plots indicate that lyxoNA, xyloNA, and DxyloNA are flexible (in that order) in comparison to corresponding C3’ inverted counterparts arabinoNA, RNA and DNA, respectively. The legends in both the plots are same.

The conformational space of the double-stranded nucleic acids was explored by calculating free energy surfaces using two reaction coordinates, viz., average twist angle (χ), and end-to-end distance (*l*) and the resulting free energy surface contours are depicted in Figure 4. The lower energy structure(s) corresponding to each of the doublestranded nucleic acids are marked and are depicted in Figure 5. Further, comparison of the free energy surfaces shown in Figure 4 and the twist angle distribution plots suggest that the sampling of the structural landscape by constraint free MD simulations is reasonable except in the case of lyxoNA. The free energy surfaces also point out that the right-handed helix of double-stranded DNA is energetically most stable while doublestranded xyloNA is least stable and the energy ordering is DNA >> lyxoNA > RNA ~ ANA >> DxyloNA > xyloNA. The RNA (R1), arabinoNA (A1) and DNA (D1) show a single minimum on the free energy surface which correspond to a right-handed double helical minimum on the free energy surface which correspond to a right-handed double helical structure. On the other hand, various minima observed on the free energy surface of xyloNA (X1, X2, X3 and X4), lyxoNA (L1), and DxyloNA (dX1 and dX2) loose helical integrity. Interestingly, lyxoNA (L1) shows a single minimum free energy circular-ladder structure (see Figure 5), with absence of helical content. Moreover, the L1 conformation shows considerable amount of base flipping, indicating the fragility of canonical Watson-Crick base paring. The xyloNA and DxyloNA show multiple minima on the free energy surface, which can also be characterised as a combination of left-handed helices and ladder structures. In general, the right-handed double helical structural integrity is compromised with the inversion of configuration of the furanose ring at the C3’ position relative ribose/arabinose.

**Figure 4.**
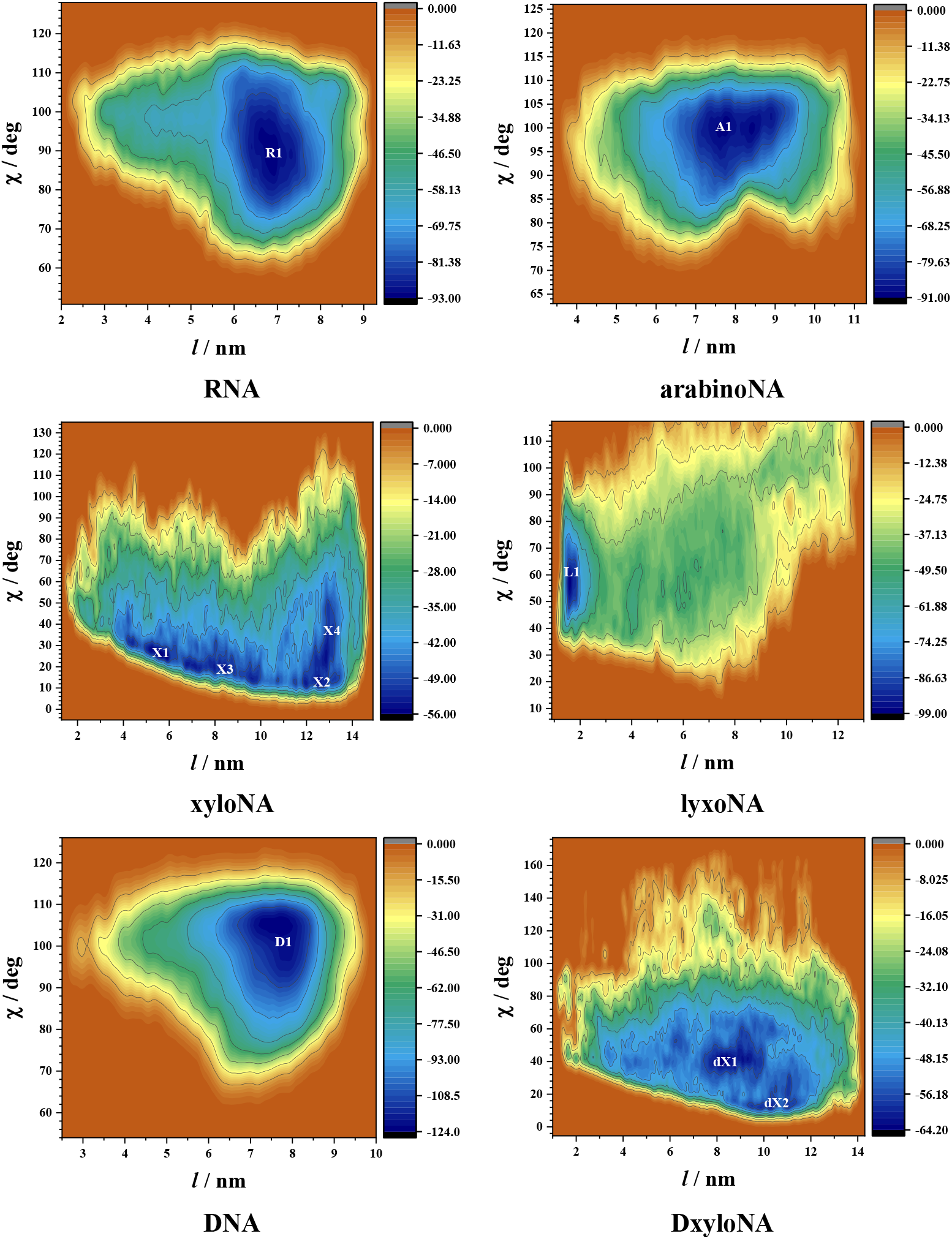
Free energy surface contours as a function of average twist angle (*χ*) and end-to-end distance (*l*) for various double-stranded nucleic acids. The corresponding low energy structures are alpha-numerically marked and are shown in Figure 5.

**Figure 5.**
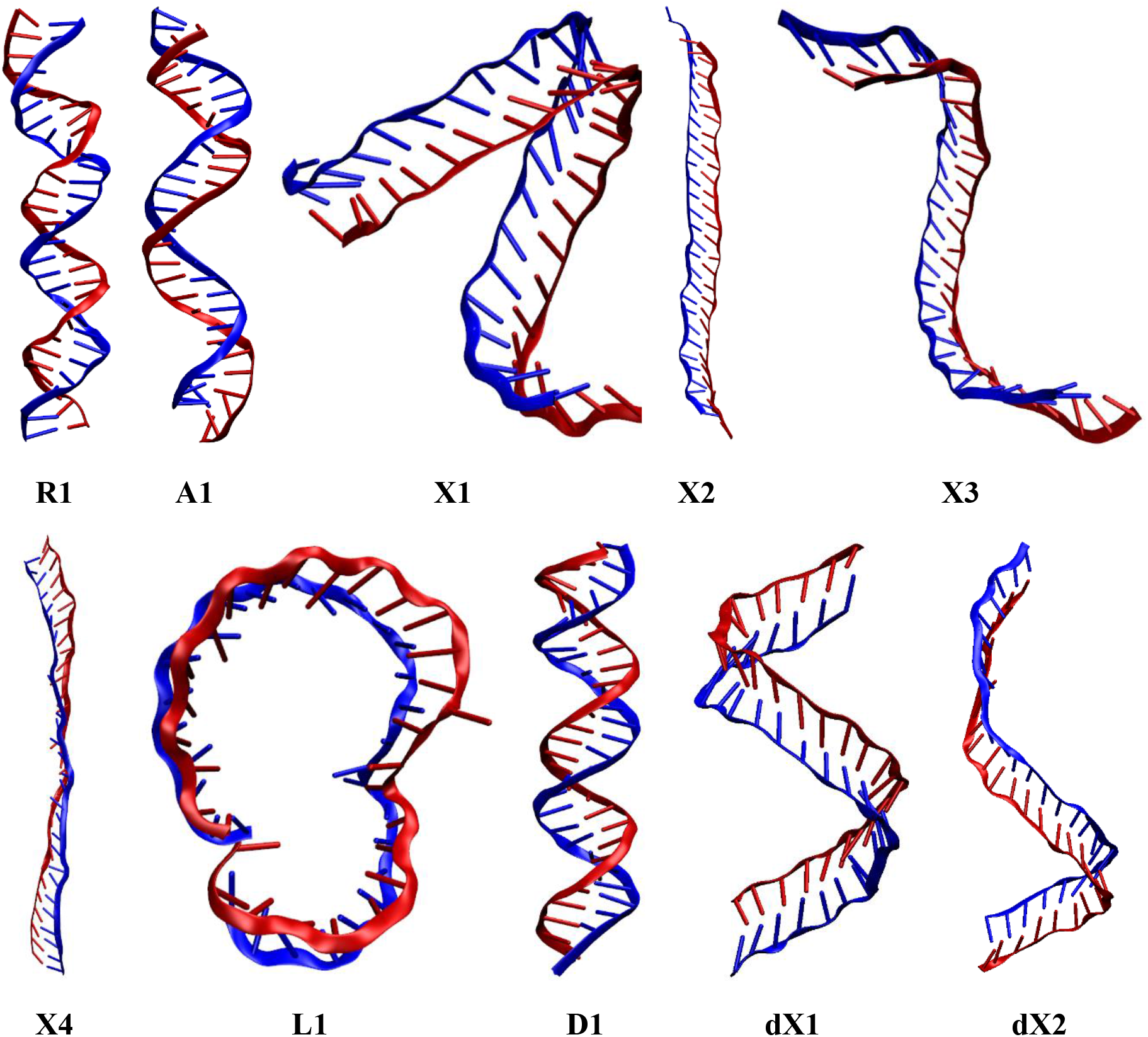
Structures of minimum free energy conformations of RNA (R1), arabinoNA (A1), xyloNA (X1, X2, X3 and X4), lyxoNA (L1), DNA (D1) and DxyloNA (dX1 and dX2) marked in the free energy surface depicted in Figure 4.

Further, various average structural parameters such as inter-strand N–N, interstrand P–P and intra-strand stacking distances were analysed and their distribution plots (averaged over five 100 ns trajectories) are shown in Figure 6. On the first glance, the distribution plots, shown in Figure 6, indicate that lyxoNA is structurally very different than rest. Additionally, the inter-strand N–N distance shows a very similar distribution for all the double-stranded nucleic acids, except in the case of lyxoNA. Similar, observations can be made from the inter-strand P–P distance distribution plot. In the case of lyxoNA increase in the N–N distance distribution and concomitant decrease in the P–P distance distribution is attributed to base-flipping, as seen in Figure 5 (see L1 structure). Furthermore, the distribution plots of intra-strand stacking distance suggest that DNA exhibits most efficient intra-strand π-π interaction of the adjacent bases with distribution maximum around 0.33 nm, followed by arabinoNA and RNA. On the other hand, lyxoNA and DxyloNA show multiple mode distribution, some of which shows the presence of π-π interaction between the adjacent bases. Contrastingly, the xyloNA shows the narrowest stacking distance distribution with maximum around 0.56 nm, which is devoid of any intra-strand π-π interaction between the adjacent bases. The formation of the lowest free energy ladder structure for the xyloNA can be inferred based on complete loss of π-π interaction between the adjacent bases. The larger, yet narrow distribution of the interstrand stacking distance can also be attributed to the formation of favourable Lpoooπ interactions between the oxygen atom of the xylose and the adjacent nucleobase due to the disposition of the sugar moiety relative to the nucleobases in xyloNA.^21^

**Figure 6.**
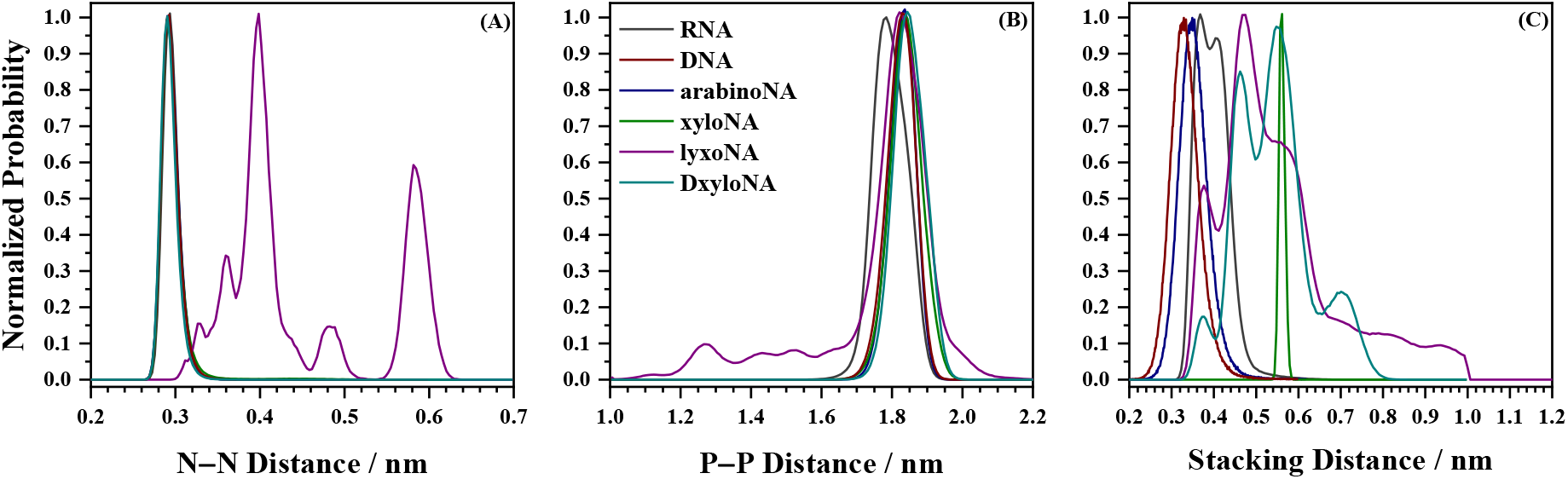
Distribution plots of (A) inter-strand N–N, (B) inter-strand P–P and (C) intrastrand stacking distances, averaged over five 100 ns trajectories for various doublestranded nucleic acids. The legends in all the three plots are same.

The presence of helicity or lack of it was analysed using 3DNA,^37,38^ for the six variants of double-stranded nucleic acids and the important helical parameters such as H-twist, major-grove width and minor-grove width are listed in Table 1. The helical parameters suggest that average H-Twist for the DNA (32.2°) and RNA (32.4°) are almost equal (the standard deviation in RNA is higher) and the corresponding value for arabinoNA is 27.2°. With similar end-to-end distances of DNA and RNA and arabinoNA (about 7.8 nm), a lower value of H-Twist for the arabinoNA indicates untwisting of arabinoNA relative to

**TABLE 1:**
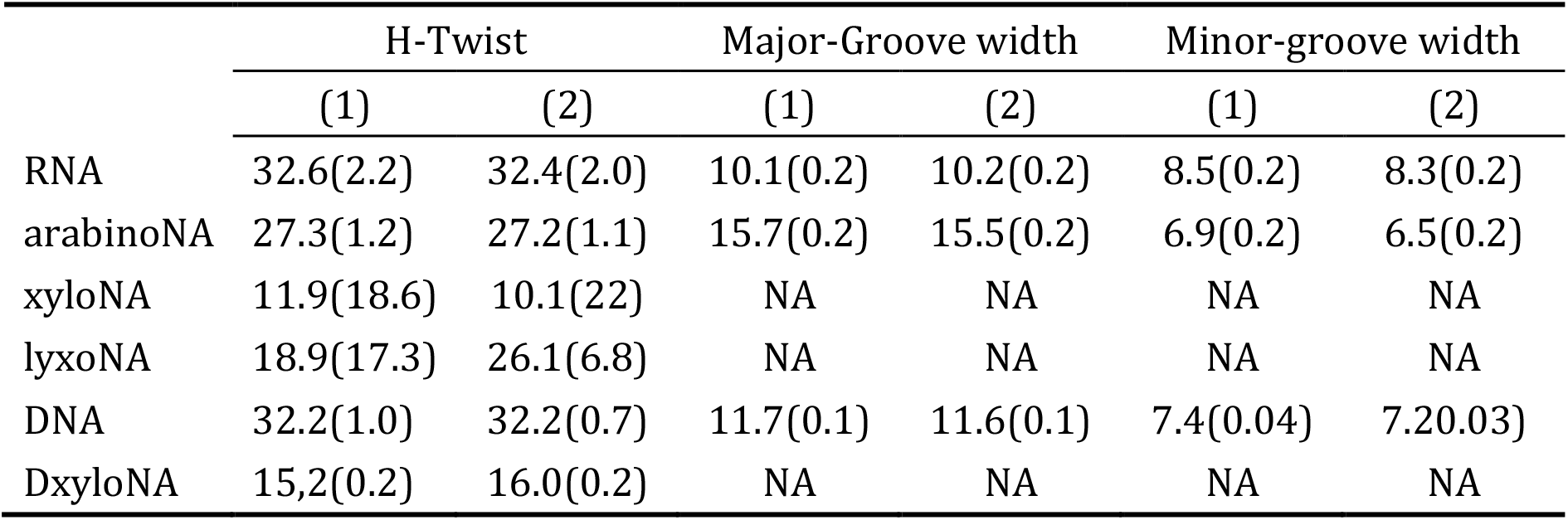
H-Twist (deg), major-grove width (Å) and minor-grove width (Å) obtained following 3DNA analysis for the double-stranded nucleic acids. In each case the first column is value obtained by averaging over all the snapshots five trajectories and the second column is the value obtained by averaging over the stable geometries depicted in Figure 5. The values in parenthesis are standard deviations.

DNA and RNA. This untwisting of arabinoNA double helix results in substantial increase in the major groove width accompanied by a marginal decrease in minor groove width, wherein, the ratio of major-to-minor groove widths for the arabinoNA (2.27) is disproportionately large in comparison with RNA (1.19) and DNA (1.58). Further, the 3DNA suggests that RNA, arabinoNA, and DNA form right-handed helical structures, while xyloNA and DxyloNA form left-handed ladder-like structures (W-form) in some instances. The most important consequence of the inversion of configuration at the C3’ position is the loss of helicity resulting in the disappearance of both the major and the minor grooves.

The structural landscape of the furanosyl double-stranded nucleic acids is primarily governed by the configuration of the C3’ position. Inversion at C3’ position relative to ribose results in loss of helicity as in the case of xyloNA and lyxoNA. These structural changes can be associated with sugar pucker, which is influenced to a larger extent by the inversion of configuration at the C3’ position. On the other hand, inversion at C2’ position (arabinoNA) leads to under-twisted helix in comparison to RNA, However, the loss of chirality at C2’ position in DNA appears to be a better proposition than inversion, which is intriguing, and is associated with change in sugar pucker. The present set of simulations bring out that incorporation of ribose results in better defined helices among the four possible furanose variants with robust structural integrity. Based on these results it might be surmised that ribose is a natural choice over other three furanoses. The formation of the helix and its associated structural features are utilized by a variety of biological machinery such as helicases, topoisomerases, groove binders, and several others, which could not have been possible with xylose and lyxose derived nucleic acids and to a much lesser degree of control with arabinoNA. Returning to the question raised by Eschenmoser,^5^ “why did nature choose furanosyl-RNA and not pyranosyl-RNA?” is still a matter of debate, however, among the furanoses, ribose appears to be the best natural choice.

## CONCLUSIONS

Summarizing, all-atom molecular dynamics simulations on six double-stranded nucleic acids viz., RNA, arabinoNA, xyloNA, lyxoNA, DNA, and DxyloNA suggest that the configuration of C3’ stereocentre determines the ability of double-stranded nucleic acids to form right-handed helices. The inversion of the C3’ stereocentre acts as a toggle switch for the helix to ladder structural transformation. On the other hand, the inversion of the C2’ stereocentre has a less pronounced effect, resulting in the modification of groove widths. A comparison of structural features suggests that ribose containing doublestranded nucleic acids are structurally robust and form better defined helices, which might have resulted evolutionary choice.

## Acknowledgements

RKS thanks IIT Bombay for the institute postdoctoral fellowship and at thanks University Grants Commission (UGC) for the research fellowship. Authors gratefully acknowledge the SpaceTime-2 supercomputing facility at IIT Bombay for the computing time. The support and the resources provided by ‘PARAM Brahma Facility’ under the National Supercomputing Mission, Government of India at the Indian Institute of Science Education and Research (IISER) Pune are gratefully acknowledged. The authors wish to thank Dr. Amutha Ramaswamy for making the initial structure of double-stranded xyloNA available. The authors also wish to thank Prof. R. B. Sunoj for his comments and suggestions.

## Author Contributions

The project was initiated by AK and carried out all the initial MD simulations with help from AT. RKS verified all the MD simulations and carried out metadynamics and free energy calculations. The results were jointly interpreted by all the authors and the manuscript was written by AK, RKS and GNP.

## Notes

The authors declare no competing financial interest.

